# Using Outliers in Freesurfer Segmentation Statistics to Identify Cortical Reconstruction Errors in Structural Scans

**DOI:** 10.1101/176818

**Authors:** Abigail B. Waters, Ryan A. Mace, Kayle S. Sawyer, David A. Gansler

**Affiliations:** Department of Psychology, Suffolk University, 73 Tremont Street, Boston, MA, USA; Department of Anatomy & Neurobiology, Boston University School of Medicine, Boston, MA, USA; VA Boston Healthcare System, Boston, MA, USA; Athinoula A. Martinos Center for Biomedical Imaging, Massachusetts General Hospital, Harvard Medical School, Charlestown, MA, USA; Sawyer Scientific, LLC, Boston, MA, USA

**Keywords:** quality assurance, automated segmentation statistics, reconstruction error, Freesurfer

## Abstract

**Introduction:** Quality assurance (QA) is vital for ensuring the integrity of processed neuroimaging data for use in clinical neurosciences research. Manual QA (visual inspection) of processed brains for cortical surface reconstruction errors is resource-intensive, particularly with large datasets. Several semi-automated QA tools use quantitative detection of subjects for editing based on outlier brain regions. There were two project goals: (1) evaluate the adequacy of a statistical QA method relative to visual inspection, and (2) examine whether error identification and correction significantly impacts estimation of cortical parameters and established brain-behavior relationships.

**Methods:** T1 MPRAGE images (N = 530) of healthy adults were obtained from the NKI-Rockland Sample and reconstructed using Freesurfer 5.3. Visual inspection of T1 images was conducted for: (1) participants (*n* = 110) with outlier values (*z* scores ± 3 *SD*) for subcortical and cortical segmentation volumes (outlier group), and (2) a random sample of remaining participants (*n* = 110) with segmentation values that did not meet the outlier criterion (nonoutlier group).

**Results:** The outlier group had 21% more participants with visual inspection-identified errors than participants in the non-outlier group, with a medium effect size (*Φ* = 0.22). Nevertheless, a considerable portion of images with errors of cortical extension were found in the non-outlier group (41%). Sex significantly predicted error rate; men were 2.8 times more likely to have errors than women. Although nine brain regions significantly changed size from pre-to post-editing (with effect sizes ranging from 0.26 to 0.59), editing did not substantially change the correlations of neurocognitive tasks and brain volumes (*ps* > 0.05).

**Conclusions:** Statistically-based QA, although less resource intensive, is not accurate enough to supplant visual inspection. We discuss practical implications of our findings to guide resource allocation decisions for image processing.

## 1. Introduction

Accurate brain volume estimation is considered essential to research brain-behavior relationships. Structural neuroimaging studies have found significant associations between regional brain volumes and domains of cognitive functioning, including executive functioning (Yuan & Raz, 2014), attention (Seidman, Valera, & Makris, 2005), and memory (Van Pettan, 2004). As neuroimaging increasingly emphasizes the use of large-scale datasets to assess these relationships, ensuring the integrity and validity of data via quality assurance (QA; also called quality control), is critical and must be done efficiently. There is an emergent need to understand the effectiveness of QA methodology to help researchers make resource allocation decisions for image processing.

Freesurfer (http://surfer.nmr.mgh.harvard.edu/fswiki) is a commonly used open source software suite for automated processing of magnetic resonance imaging (MRI) data. Although validation studies have shown that automated segmentation in Freesurfer is commensurate to manual measurement (Fischl et al., 2002), post-reconstruction visual inspection of cortical “pial” surface segmentation has become common practice to identify incorrect inclusion of non-brain tissues (Desikan et al., 2010). Manual QA (i.e., visual inspection of imaging slices) used to confirm the validity of reconstructed images is time-and resource-intensive, particularly for large-scale neuroimaging datasets. QA methods vary greatly between studies (e.g. Chen et al., 2015; Kaufmann et al., 2017, Ahmed et al., 2015); some studies use manual inspection, semiautomated QA, or a combination of both methods. This variation may differentially influence neuroimaging data used in research analyses.

Semi-automated QA methods for reconstructed images are needed to address limited resources in neuroimaging research. Both the QA toolkit in the Freesurfer software suite (Koh, Lee, Pacheco, Pappu, & Vinke, 2017) and the Mindcontrol web application (Keshavan et al., 2017) use statistical analyses of cortical and subcortical regions to identify images that may need manual correction. These regional volumetric measurements are automatically generated based on the probabilistic location of structures (Fischl et al., 2002; Fischl et al., 2004), which due to individual variation of anatomical features, may result in errors of cortical inclusion and exclusion. Therefore, regions with surface reconstruction errors would have over-or underestimation of volumetric measurement resulting in statistical outliers.

The primary objective of this study was to investigate whether anomalies in cortical and subcortical volumetric measurements are associated with reconstruction errors, thereby testing the assumption that statistically-based methods can identify images with reconstruction errors. In addition, we investigated whether participant characteristics (age and sex) impacted the odds of reconstruction errors. This study focused on errors where the cortical surface extended into nonbrain tissues. Manual correction of the white matter surface using control points does not result in significantly different volumetric measurements (McCarthy et al., 2015); therefore, errors in delineation of white matter are unlikely to result in statistical outliers. We hypothesized that the outlier images would have significantly more cortical surface errors than those not identified via this method. We did not have a priori hypotheses regarding the influence of participant characteristics.

A secondary objective was to determine whether the identification and correction of cortical boundary errors significantly impact established brain-behavior relationships. We investigated whether the association between neurocognitive measures and brain volumes would significantly differ from pre-to post-editing. We expected that some volumes (i.e., precentral, postcentral gyrus) would be more affected by errors than others, given previous literature (Keshavan et al., 2017). We had no a priori hypothesis regarding the impact of editing on brain-behavior relationships. We discuss the implications of our findings in the context of guiding resource allocation decisions regarding neuroimaging QA and the potential influence on the analysis of brain-behavior relationships.

## 2. Methods

### 2.1 Participants

De-identified phenotypic and neuroimaging data for 645 participants was made available via the enhanced Nathan Kline Institute – Rockland Sample (NKI-RS), an open-access, cross sectional, community sample (Nooner, et al., 2012). Rockland County’s economic and ethnic demographics are representative of the United States census (U.S. Census Bureau, 2009), making the NKI-RS generalizable to the U.S. population. Zip-code based recruitment was conducted via mailings, flyers, and electronic advertisement. Data use agreement was accepted by NKI-RS and data handling procedures were approved by the Institutional Review Board at Suffolk University.

The NKI-RS excludes participants if they were diagnosed with or reported severe psychiatric disorders (bipolar disorder, schizophrenia disorder, schizoaffective disorder), severe developmental disorders (autism spectrum disorders, intellectual disabilities), current suicidal or homicidal ideation, severe cerebral trauma (stroke, moderate to severe traumatic brain injury, transient ischemic attack in the past two years), severe neurodegenerative disorders (Parkinson’s disease, Huntington’s disease, dementia), a history of substance dependence in the past two years (except cannabis), a lifetime history of psychiatric hospitalization, current pregnancy, or MRI contraindications. Participants were excluded from these analyses if T1-weighted structural images were incomplete or had significant image artifacts. A total of 530 adult participants from the NKI-RS who met eligibility criteria and had complete scan data were included in this study (Figure 1 describes participant flow for study sample).

**Figure 1.**
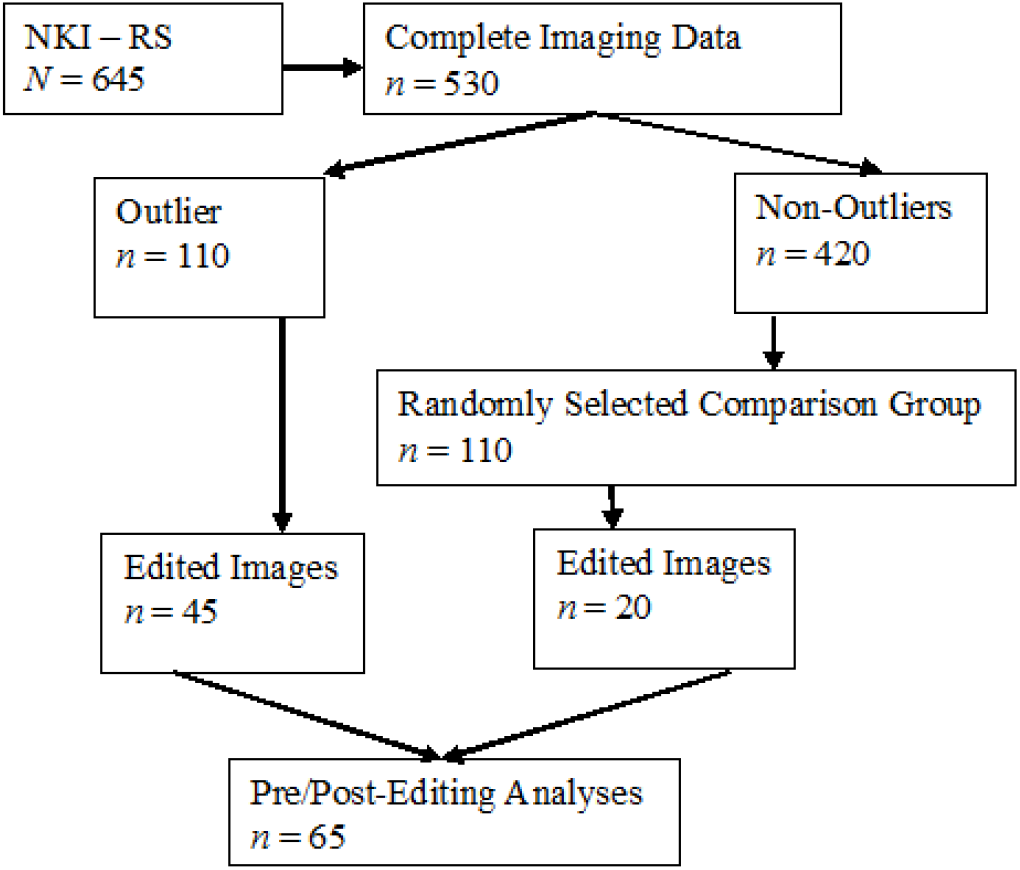
Flowchart of Nathan Kline Institute – Rockland Sample (NKI-RS) participants and the study sample.

### 2.2 Imaging

Structural images were acquired using a 3T Siemens Trio scanner (T1 MPRAGE, voxel size = 1.0 x 1.0 x 1.0 mm, 176 slices, echo time = 2.52 ms, repetition time = 1900 ms, field of view = 250 mm). MPRAGE data obtained from the NKI-RS dataset are available in their raw form. DICOM data were converted to the mgz format and MPRAGE images were auto-reconstructed in Freesurfer 5.3. Each image was mapped into standard morphological space (MNI305; Collins, 1994), and volumes were generated for cortical and subcortical regions based on the Desikan-Killiany Atlas (Desikan et al., 2006), which included white matter, gray matter, and other anatomical features.

### 2.3 Outlier identification and comparison group

Two participant subsamples were identified from the 530 participants that met study inclusion criteria. First, Freesurfer-automated segmentation statistics for subcortical and cortical segmentations were standardized. Next, all participants with z-scores 3 *SD* above or below the mean for one or more normalized brain volume labels estimated by Freesurfer (66 total automated segmentation brain region variables) were identified as the *outlier group (n* = 110). This statistically-based identification method was designed to assess the underlying assumption that segmentation statistics can be used to identify incorrectly reconstructed images, which is the basis of many semi-automated QA techniques (Kaufmann et al., 2017, Koh, Lee, Pacheco, Pappu, & Vinke, 2017).

The 3 *SD* cut-off was chosen because 2 *SD* cut-off was too broad for practical reasons (it identified 311 of 530 images as outliers) and would most likely result in low specificity. If outlier measurements predict error rates, we hypothesized that the more extreme outliers would have greater specificity. Finally, a random sample of participants that did not meet the outlier criterion (i.e., no standardized segmentation brain volumes ± 3 *SD)* were selected via random number generation for the *non-outlier group (n* = 110), which served as a comparison group. Table 1 presents the demographic characteristics for both groups, which represent the final sample for data analysis (N = 220).

**Table 1.** Basic Demographics by Group

| | Outlier group ($n = 110$) | | Non-Outlier group ($n = 110$) | | Total Sample ($N = 220$) | |
| --- | --- | --- | --- | --- | --- | --- |
|  | *M* | *SD* | *M* | *SD* | *M* | *SD* |
| Age | 53.57 | 18.92 | 43.90 | 17.19 | 48.74 | 18.67 |
|  | % |  | % |  | % |  |
| Sex |  |  |  |  |  |  |
| Male | 50 |  | 29.1 |  | 39.5 |  |
| Female | 50 |  | 70.9 |  | 60.5 |  |
| Race |  |  |  |  |  |  |
| Native Am. | 0.9 |  | 0 |  | 0.5 |  |
| Asian | 3.6 |  | 6.4 |  | 5.0 |  |
| Black | 14.5 |  | 18.3 |  | 16.5 |  |
| White | 79.1 |  | 73.4 |  | 76.6 |  |
| Other | 1.8 |  | 1.8 |  | 1.4 |  |

### 2.4 Error identification

Because almost all of the identified outlier regions were ≥ 3 *SD* above the mean (only 8.2% of outliers were found at least 3 *SD* below the mean), investigators hypothesized that the statistical method was best suited to identify erroneous inclusion of skull and dura into cortical volumes. Therefore, error identification focused on errors of cortical boundary extension (See Figure 2 for several example images). All brain images were visually inspected using Freeview to identify the presence or absence of errors in reconstruction. Graduate students (A.W. and R.M.), who were trained in visual inspection by a senior author (K.S.) with extensive experience in imaging science, served as raters.

Both graduate students performed manual visual inspection on each of the auto-reconstructed brains for both outlier and non-outlier participants (N = 220) in Freeview. White and gray matter surfaces were overlaid onto T1 scans (i.e., before skull stripping) to assist in cortical identification. All slices were viewed in the coronal plane. Each image was coded as having no errors (0) or projection errors (1). A projection error occurred when the cortical boundary incorrectly extended outward more than approximately 3 voxels into the dura and/or skull on at least one slice. The trained graduate students completed their ratings independently and were blind to participant characteristics (including outlier or non-outlier group membership). Both raters recorded their viewing time using a stopwatch.

**Figure 2.**
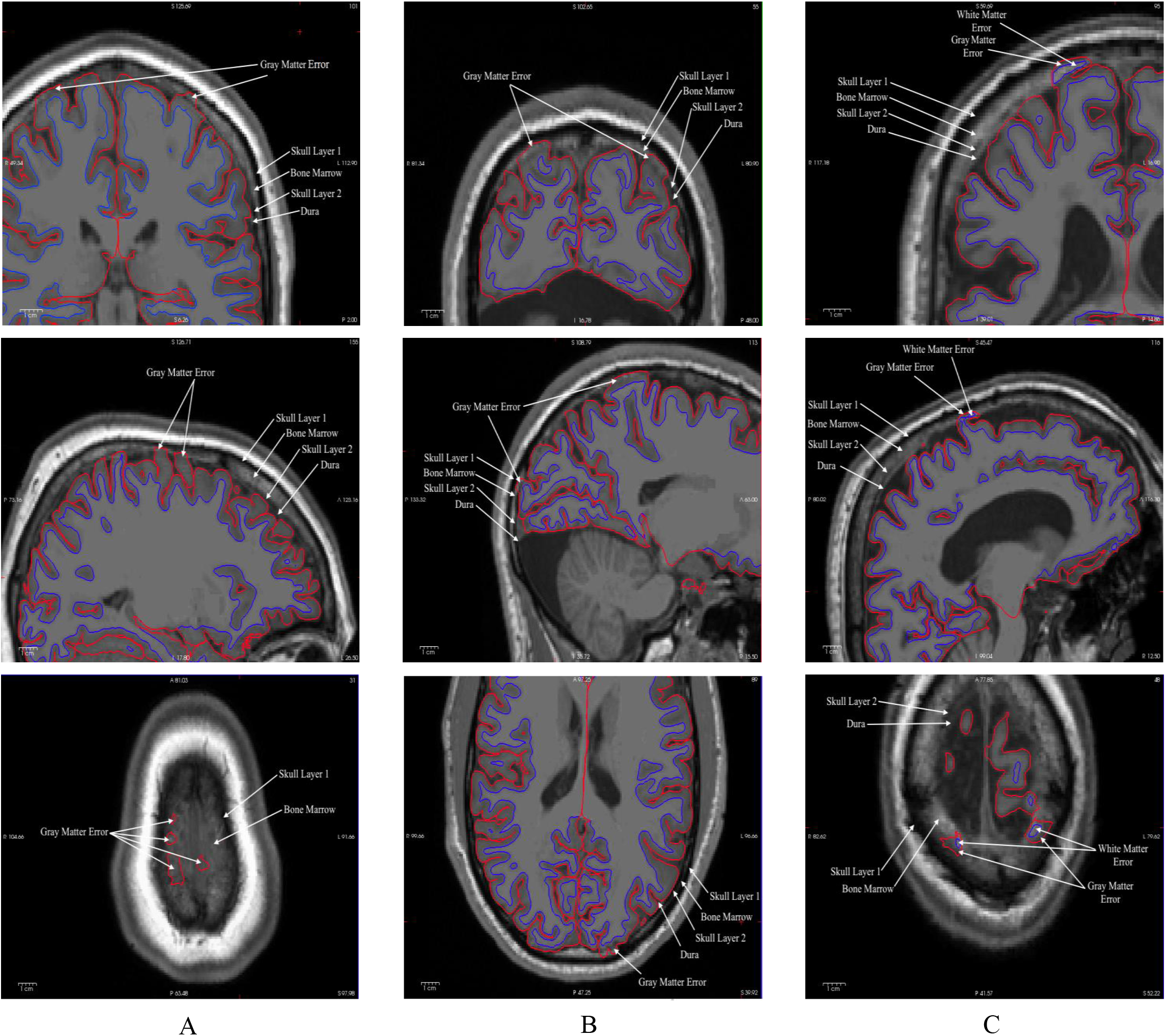
Coronal, Sagittal, and Horizontal View of (A) Errors of cortical extension in the precentral/post-central region; (B) Errors of cortical extension in posterior regions; and (C) Errors of cortical extension that include both white and grey matter.

On average, raters took 70 seconds to visually inspect and identify errors for each image. This estimate does not include the time required to load scan data in Freeview (approximately 20 seconds), which depended on computer speed. Interrater reliability analyses indicated moderate agreement (Cohen’s Kappa = 0.65, 95% CI: 0.55 – 0.74; 82.4% agreement), with 181 of the 210 ratings consistent between both raters. For the 39 discordant ratings, consensus was reached with both raters and a senior author (KS). Most discordant images (76.9%) were assigned consensus ratings of no error. Twenty-four of these 39 images were assigned a rating of no error because they did not meet full criteria for an error (> 3 voxels cortical extension), but contained some partial or full voxel cortical extension (some for large areas of the cortical surface; see Supplemental Figure 1). Due to variability in scan quality and anatomical features, raters concluded that these images could not be conclusively categorized as errors of cortical extension. These images were instead categorized as no error images. These ratings were combined with the agreement ratings (181 images) and used in all subsequent analyses. In total, 138 of the 210 images had errors (65.7%).

### 2.5 Group and error characteristics

Between group t-tests, chi-square tests, and logistic regression were used to determine if group membership (outlier versus non-outlier) was significantly associated with reconstruction error or other participant characteristics. Further analyses were performed to assess characteristics associated with errors, regardless of group membership.

### 2.6 Brain volumes and measures of neurocognitive functioning

Pearson correlations were used to test the associations between brain regions and neurocognitive tasks. Correlation coefficients were compared between images pre-and post-editing using a Fisher r-to-*z* transformation (Meng, Rosnthal, & Rubin, 1992) and by comparing effect sizes (Cohen, 1988).

A subsample of images with errors (65 of 138 images) were manually edited (Savalia, Agres, & Wig, 2015), and cortical regions were averaged across hemispheres for the purpose of data reduction. Volumes were compared before and after editing (Table 2). Nine brain volumes across all edited images showed significant decreases in volume from pre-to post-editing.

These nine volumes were used to assess brain-behavior relationships. Pearson correlation coefficients were also used to estimate the associations between edited values and measures of neurocognitive functioning to determine if error correction meaningfully influenced the outcome of brain-behavior relationships. Tasks were chosen based on previous literature to assess the most robust brain-behavior relationships (Yuan & Raz, 2014; Phan, Wager, Taylor, & Liberzon, 2002; Ward & Frackowiak, 2003).

**Table 2.** Mean Decreases in Volumes (mm^3^) Pre-to Post-Editing (N = 65)

|  | *M* | *SD* | 95% CI |  | *t* | *p* | *d* |
| --- | --- | --- | --- | --- | --- | --- | --- |
|  |  |  | Lower | Upper |  |  |  |
| Precentral | 205.33 | 345.79 | 119.65 | 291.01 | 4.79 | <0.001 | 0.59 |
| Postcentral | 221.45 | 461.53 | 107.09 | 335.82 | 3.87 | <0.001 | 0.48 |
| Superior Parietal | 117.82 | 281.35 | 48.10 | 187.53 | 3.38 | 0.001 | 0.42 |
| Posterior Cingulate | 12.33 | 32.85 | 4.19 | 20.47 | 3.03 | 0.004 | 0.38 |
| Inferior Parietal | 132.58 | 386.33 | 36.85 | 228.30 | 2.77 | 0.007 | 0.34 |
| Isthmus Cingulate | 10.80 | 37.32 | 1.55 | 20.05 | 2.33 | 0.023 | 0.29 |
| Precuneus | 12.15 | 46.96 | 0.52 | 23.79 | 2.09 | 0.041 | 0.26 |
| Superior Frontal | 57.04 | 220.52 | 2.40 | 111.68 | 2.09 | 0.041 | 0.26 |
| Caudal Middle Frontal | 35.18 | 137.32 | 1.16 | 69.21 | 2.07 | 0.043 | 0.26 |
| Fusiform | 32.55 | 147.25 | -3.93 | 69.04 | 1.78 | 0.079 | 0.22 |
| Rostral Middle Frontal | 50.85 | 240.63 | -8.78 | 110.47 | 1.70 | 0.093 | 0.21 |
| Parsorbitalis | 5.97 | 28.50 | -1.09 | 13.03 | 1.69 | 0.096 | 0.21 |
| Rostral Anterior Cingulate | 10.11 | 48.84 | -1.99 | 22.21 | 1.67 | 0.100 | 0.21 |
| Caudal Anterior Cingulate | 6.65 | 32.81 | -1.48 | 14.78 | 1.63 | 0.107 | 0.20 |
| Inferior Temporal | 49.22 | 249.59 | -12.62 | 111.07 | 1.59 | 0.117 | 0.20 |
| Lingual | 28.65 | 146.77 | -65.02 | 7.72 | -1.57 | 0.121 | 0.20 |
| Supramarginal | 96.60 | 496.99 | -26.55 | 219.75 | 1.57 | 0.122 | 0.19 |
| Medial Orbitofrontal | 15.61 | 82.56 | -4.85 | 36.07 | 1.52 | 0.132 | 0.19 |
| Parstriangularis | 13.60 | 73.57 | -4.63 | 31.83 | 1.49 | 0.141 | 0.18 |
| Paracentral | 6.96 | 44.67 | -4.11 | 18.03 | 1.26 | 0.214 | 0.16 |
| Middle Temporal | 39.75 | 276.40 | -28.73 | 108.24 | 1.16 | 0.251 | 0.14 |
| Parahippocampal | 6.92 | 53.18 | -6.26 | 20.09 | 1.05 | 0.298 | 0.13 |
| Temporal Pole | 6.86 | 54.70 | -6.69 | 20.42 | 1.01 | 0.316 | 0.13 |
| Insula | 8.93 | 89.67 | -13.29 | 31.15 | 0.80 | 0.425 | 0.10 |
| Pericalcarine | 2.97 | 31.66 | -4.88 | 10.81 | 0.76 | 0.452 | 0.09 |
| Bankssts | 3.61 | 47.20 | -8.09 | 15.30 | 0.62 | 0.540 | 0.08 |
| Frontal Pole | 2.09 | 27.83 | -8.99 | 4.80 | -0.61 | 0.547 | 0.08 |
| Parsopercularis | 4.12 | 69.96 | -13.22 | 21.45 | 0.47 | 0.637 | 0.06 |
| Lateral Orbitofrontal | 4.92 | 89.43 | -27.08 | 17.24 | -0.44 | 0.659 | 0.06 |
| Superior Temporal | 8.37 | 188.52 | -38.34 | 55.08 | 0.36 | 0.722 | 0.04 |
| Transversetemporal | 0.33 | 12.04 | -2.65 | 3.31 | 0.22 | 0.825 | 0.03 |
| Lateral Occipital | 3.73 | 153.26 | -41.71 | 34.24 | -0.20 | 0.845 | 0.02 |
| Cuneus | 0.89 | 46.45 | -12.40 | 10.62 | -0.16 | 0.877 | 0.02 |
| Entorhinal | 0.97 | 72.06 | -16.89 | 18.83 | 0.11 | 0.914 | 0.01 |

3 *Note. Volumes in descending order by Cohen's d effect size.*

Participants included in these analyses had completed the Delis-Kaplan Executive Functioning System (D-KEFS), the Wechsler Abbreviated Scale of Intelligence (WASI-II), and the Penn Computerized Neuropsychological Test Battery (Penn CNB) during their involvement in NKI-RS. All tests were administered per the guidelines presented in their respective administration manuals by trained research assistants. All tests on the Penn CNB were part of a larger computerized battery that participants completed on a desktop with trained research assistant supervision.

#### 2.6.1 D-KEFS Trails

The Trails subtest on the D-KEFS includes five conditions. Completion time in seconds from the first, fourth, and fifth conditions were included in analyses. These conditions assess visual attention, mental flexibility, and visuo-motor coordination, respectively (Kaplan, Delis, & Kramer, 2001).

#### 2.6.2 WASI Perceptual Reasoning Index (PRI)

The WASI-II PRI was calculated using the standard scores from two subtests: Block Design and Matrix Reasoning. The Block Design subtest assesses spatial visualization, abstract conceptualization, and visuo-motor coordination. The Matrix Reasoning subtest assesses non-verbal fluid reasoning (Weschsler, 2011).

#### 2.6.3 Penn CNB Conditional Exclusion Task (PCET)

The PCET was developed to assess mental flexibility and abstraction (Gur et al., 2001).

#### 2.6.4 Penn CNB Emotion Recognition Task (ERT)

The ERT is a measure of emotion recognition along a continuum of expression intensity (Gur et al., 2001).

## 3. Results

### 3.1 Error characteristics and statistical method

The rate of reconstruction errors was 62.7% in the outlier group and 40.9% in the nonoutlier group (Table 3). There were significant differences in error rates between the non-outlier and outlier groups, *(X^2^* (1, 220) = 10.49, *p* = 0.001), with a medium effect size (Φ = 0.22). Based on the odds ratio, the outlier images were 1.69 times more likely to have a projection error than the randomly selected images. Raters identified 45 out of 220 images (20.5%) in need of further reconstruction that were not detected by the statistical method (i.e., false negative). The statistical method had an accuracy of 60.9%, a sensitivity of 60.5%, and a specificity of 59.1%.

**Table 3.** Crosstabulation of Group Status and Error Rate

|  | Outlier Group<br>(Predicted Positive) | Non-Outlier Group<br>(Predicted Negative) | Total |
| --- | --- | --- | --- |
| Error<br>(Positive) | 69<br>(62.7%; 60.5%) | 45<br>(40.9%; 39.5%) | 114<br>(51.8%) |
| No Error<br>(Negative) | 41<br>(37.3%; 38.7%) | 65<br>(29.5%; 61.3%) | 106<br>(48.2%) |
| Total | 110<br>(50%) | 110<br>(50%) | 220 |

*Note.* $*p \leq .001$ . Numbers in parentheses indicate: (column %; row %). There were significant differences in error rates between the non-outlier and outlier groups, ( $X^2 (1, 220) = 10.49, p = 0.001$ ), with a medium effect size ( $\Phi = 0.22$ ).

Outliers were found in 55 of 66 segmentation volumes. Outliers in the ventricles and surface hole regions were most frequent, as 56.4% of images in the outlier group were identified using statistical outliers in the ventricular, surface hole, vessel, and CSF volumes alone (13 of 66 brain region). To verify that ± 3 *SD* criterion used in outlier identification was not overly stringent, chi-square tests were performed on the 220 error-rated images using the less conservative 2.5 *SD* and 2 *SD* cut-offs. Both the 2.5 *SD* cut-off (X^2^ (1, 220) = 10.49, *p* = 0.001) and the 2 *SD* cut-off (X^2^ (1, 220) = 10.49, *p* = 0.001) showed significant differences in error rates between outlier images and images that did not meet cut-off criteria. However, the proportion of images with errors that did not meet cut-off criterion remained relatively consistent (41.4% and 40% respectively) as stringency was reduced. The proportion of images with errors for the outlier group decreased (60.3% and 56.3% respectively), even as the group size expanded to 121 for 2.5 *SD* and 160 for 2 *SD*. Reducing the stringency did not improve sensitivity or specificity of the statistical method.

Age (*t*(218)= -3.97, *p* < .001) and sex *(X^2^* (1, 220) = 10.06, *p* = 0.002) were significantly different between the two groups, with the outlier group 9.7 years older on average and 21% more male. A series of logistic regression analyses were used to examine whether participant characteristics were associated with error rate across groups. Age and sex were entered into the model as predictor variables for projection errors (criterion). The model was statistically significant and explained an estimated 6% (Cox and Snell *R^2^)* to 9% (Nagelkerke *R^2^)* of the variance in projection errors (X^2^ = 14.91, *p* = 0.001). Sex was the only significant predictor of error rate (67.8% of men had errors, compared to 41.6% of women), with women being 66% less likely to have projection errors than men *(OR* = .34, 95% CI = .19 - .59).

### 3.2 Brain volumes and measures of neurocognitive functioning

A Kruskal-Wallis one-way analysis of variance did not yield significant gender or group effects on error size, as measured by changes in volume post-editing (i.e., the ranks were unchanged by editing). Therefore, all edited images were analyzed together across groups to assess brain-behavior relationships. Of the 34 brain regions, nine (26.5%) showed significant decreases in volume from pre-to post-editing: the inferior parietal, superior parietal, posterior cingulate, isthmus cingulate, precuneus, precentral, postcentral, superior frontal and caudal middle frontal volumes (See Table 2). Cohen’s *d* ranged from 0.26 to 0.59, with all but one meeting criteria for a small effect size (0.20 < *d* < 0.50). None of the associations between selected ROIs and the neurocognitive tasks significantly changed from pre-to post-editing, as estimated by the Fisher z transformation. Pre-editing Pearson’s r-values ranged from -0.35 to 0.24, while post-editing r-values ranged from -0.35 to 0.23; all correlations meeting Cohen’s *d* criteria for a small to no effect size. The change in *r* from pre-to post-editing ranged from -.02 to .08. Of 54 correlations, one resulted in changes of effect size categorization: PCET and the isthmus cingulate volume met Cohen’s *d* criteria for a small effect size before editing (r = .20) and did not meet criteria after editing (r = .19). However, the pre-and post-editing correlations between PCET and the isthmus cingulate volume did not approach significance.

## 4. Discussion

The principal findings were that: (1) cortical reconstruction error rates were higher in a group identified by a statistical outlier QA method; (2) reconstruction error rates were too prevalent in a randomly identified non-outlier group to conclude that identification by the statistical outlier method alone was effective; and (3) while post-editing volume estimations were significantly lower in a number of instances, these putatively more accurate volume estimations did not meaningfully impact the outcome of a brain-behavior analysis. To our knowledge, this is the first study investigating the effects of QA error correction on associations between brain and neurocognitive measures. Contrary to our hypothesis that errors would only affect surface structures, we found significant decreases in both lateral and midline structures from pre-to post-editing. Regarding participant characteristics, we found that while both age and sex were associated with statistical outlier status. Only sex was significantly associated with error occurrence.

Although the use of T2-weighted scans in conjunction with T1 MPRAGE improve segmentation accuracy (Lichy et al, 2005; available with FreeSurfer v5.3 and above), there is still a need for consistent and efficient QA methodology for reconstruction of T1 scans alone. Previous research on statistically-based QA of automated reconstruction has noted the necessity of establishing a link between summary statistics and cortical extension errors. Similarities in signal intensity between cortex and dura on T1 MPRAGE scans make distinguishing the two difficult, even when using visual inspection (Viviani et al., 2017). This difficulty was apparent during this study, as trained graduate student raters with comparable experience had only “adequate” interrater reliability. Given the variability in QA methodology, statistically based QA presents the potential for a more standardized approach (Keshavan et al., 2017).

However, our use of statistical outliers in segmentation statistics to identify images with reconstruction errors was insufficient to identify all images with errors. Given the frequency of errors in the non-outlier group, one could expect approximately 40% of errors to go unidentified. These missed errors would be biased toward younger, female brains. Contrary to our hypothesis, statistically-based identification did not reliably identify inflated values resulting from cortical extension errors. Instead, images in the outlier groups are most often identified by inflated ventricle size or other measures of atrophy. These measures of atrophy are most likely age-related, given the significantly higher age in the outlier group. Perhaps statistically based methods are not identifying errors themselves, but rather anatomical features associated with poor registration and reconstruction in the Freesurfer automated pipeline. Poor registration may also account for the higher proportion of errors among male brains, given that men generally have larger ventricular volumes (Lenroot et al., 2007). It is possible that regression based QA diagnostics tailored by subject variables (e.g. age, gender) could increase the error detection rate. It is unclear from our analysis if male brains have a higher incidence of errors because of poor registration. Alternatively, the statistical method may be confounded by unknown additional factors that account for both error rates and participant characteristics.

Additionally, we observed a disjunction between the criterion stringency and the detection of errors. Had the outliers been related to the errors themselves, the proportion of errors would increase as stringency decreased. The degree to which atypical anatomical features contribute to poor registration, and by extension errors in reconstruction, seems to be restricted to the most extreme outliers. Refining the statistical criterion did not increase the specificity or sensitivity of identifying statistical outliers with existing methods. Statistically based methods were biased toward brains with age-related atrophy, as indicated by the outlier frequency among measures of atrophy and the age differences between groups. Studies that use this method alone to identify images for surface editing may introduce confounding variables into their data. This may be especially salient for research in aging or degenerative diseases, but further research is warranted.

Our findings regarding the precentral and postcentral gyri are consistent with the patterns of cortical misclassification found during the development of Mindcontrol (Keshavan et al., 2017), where these regions were identified as being more prone to errors of cortical extension. We identified additional cortical areas where errors are more likely to affect the volumetric estimation. Editing resulted in significant decreases of volume in broader aspects of the parietal, frontal, and cingulate regions. Although the focus of this study was on errors of cortical extension (i.e., errors in the boundary between gray matter, dura, and skull), mid-line structures (i.e., posterior and isthmus cingulate) significantly changed from pre-to post-editing. We found several errors of cortical extension during visual inspection that included both gray and white matter where bone marrow was included in white matter estimates (Figure 1). Because of the intensity of bone marrow, Freesurfer was more likely to characterize these as white matter and adjust the gray matter boundary accordingly.

Despite the limitations of the statistically based QA, manual inspection may be too resource intensive for large datasets. For the NKI Rockland Sample of 530 subjects, it would have taken approximately 10.5 hours to identify all images that require editing to correct pial surfaces. Accounting for the time it took for raters to load the image (approximately 20 seconds), it would have taken 13.25 hours. This does not include the considerable time (approximately 1 to 2 hours) it would have taken to edit these images, which we would estimate to be approximately 50% of the sample given error rates of scan reconstructions in this study, for a total time of approximately 411 hours for 530 scans.

Directing QA efforts toward brain regions most commonly affected by reconstruction error may lessen the resource commitment to manual inspection. Although errors may be present in other brain regions, they are unlikely to result in significant changes pre-to post-editing. This trade-off between time and likelihood of error occurrence can be considered when prioritizing error correction efforts. While there were statistically significant decreases in some brain regions post-editing (26.5%), editing resulted in practically insubstantial decreases in volume (0.1% to 2.3%). This is consistent with previous research on the effects of control points to manually edit segmentation between white and gray matter (McCarthy et al., 2015). This suggests that manual intervention does not produce incremental utility for cortical surface boundaries either.

Editing images did not significantly impact the relationship between brain volumes and neurocognitive measures. Based on our findings, we expect that allowing these errors to exist as noise in the dataset may decrease statistical power but not confound results. Although error occurrence is more likely in male brains, errors do not significantly impact associations with neurocognitive variables and usually do not significantly change volumetric estimations. Depending on the goals of individual studies and availability of resources for QA, researchers may find that the costs of visual inspection outweigh potential benefits of manual intervention in large datasets. Further research is needed to determine whether these results are replicable for other brain-behavior relationships, clinical samples, or for other imaging techniques (i.e., fMRI, PET).

There were several limitations in this study. Interrater reliability for error identification was only at the adequate level. Although consensus ratings were reached, this finding reflects the subjective nature of error identification in QA. There is no consensus regarding the threshold of error size for determining images that need manual edits. Error identification is further complicated by variation in anatomical features and scan quality. Viewing surfaces overlaid onto T1 images before skull stripping was essential to identifying errors and should be incorporated into QA processes whenever feasible.

Errors were defined in this study as extensions of three or more voxels outside the cortical boundary to limit the effects of variability in scan quality that made the barrier between dura and cortex unclear. There were 24 images which had partial or full voxel extensions (typically 1 voxel) that did not meet criteria for an error (Supplemental Figure 1). These extensions often continued across multiple slides over the top of the cortex. Further research is needed to explore the effect of small but pervasive extensions of cortex.

This study utilized open-access structural data from healthy, community dwelling older adults. Given the bias in identification toward images with inflated measures of atrophy, these findings cannot be generalized to clinical populations with increased cortical deterioration. Future research is needed to examine the impact of pathological structural changes on quality of reconstruction and error rates. Additionally, identification of specific anatomical features associated with poor reconstruction may help to increase sensitivity and specificity for statistically based QA. Future studies could use a receiver operator characteristic (ROC) curve analysis to explore cutoffs, based on a balance of sensitivity and specificity, for number of total outliers in identifying images with reconstruction error. Given the sex bias found in our studies, ROC analyses should be conducted for men and women separately.

## Conclusions

This study highlights the limited incremental utility of correcting errors of cortical extension to assess the relationships between brain volumes and neurocognitive measures. Utilizing statistically based methods alone can introduce confounds by differentially identifying older, male brains for editing. This finding is especially important for researchers utilizing large-scale datasets, given the resource commitment to manual QA intervention.

## Acknowledgements

The authors would like to acknowledge the following people and organizations for their contributions:

Douglas Greve at the MGH/HST Athinoula A. Martinos Center for Biomedical Imaging for his comments and consultation.

The NKI-Rockland Sample Initiative for providing the data used in these analyses (data collection funded through NIMH BRAINS R01MH094639-01).

The Suffolk University Psychology Department for their support of doctoral students and David Gansler’s Lab, and the contributions of undergraduate students Ms. Paige Kawai and Ms. Leah Pedersen.

